# Measurements of EEG Alpha Peak Frequencies Over the Lifespan: Validating Target Ranges on an In-Clinic Platform

**DOI:** 10.1101/2021.10.06.463353

**Authors:** David Joffe, D S Oakley, F. Arese Lucini, F. X. Palermo

**Affiliations:** WAVi Research, Boulder CO; University of Colorado School of Medicine, Aurora, CO

**Keywords:** Electroencephalogram (EEG), Alpha Power, Alpha Frequency, Alpha Peak Frequency (AF), Peak Individual Alpha Frequency (IAF), P300, Event Related Potential (ERP), Brainwave, Amplitude, Trail Making Test, Physical Reaction Time

## Abstract

**Background:** Peak individual alpha frequencies (IAF) extracted from an EEG exam can provide novel sources of information regarding brain function. This information can help measure and track changes in cognition arising from conditions such as concussion or unhealthy aging.

**Objective 1:** To validate a method for combining eyes-closed EEG with eyes-closed audio P300 ERP in order to streamline testing times involved in IAF extraction.

**Objective 2:** To validate age-stratified target ranges of IAF collected in clinic against published research

**Objective 3:** To validate the stability of IAF for data collected in routine clinical settings.

**Participants:** Two thousand twenty-five subjects aged 13-90.

**Methods:** EEG with audio P300 was collected as part of a health screening exam for studies through Colorado University, Children’s Hospital Colorado, Boone Heart Institute, WAVi Co., and various clinics alongside other clinical evaluations.

**Results:** (1) No differences were seen between IAF extracted during an eyes-closed resting and the P300 protocol. (2) The age-related AF trends measured in clinic match the age-related trends from previous research. (3) IAF remained stable over the course of 0-2 years in a test-retest dataset.

**Conclusion:** In-clinic measures of peak EEG frequency corroborate the age-related trends of published research taken over the last several decades and IAF. These results also confirm that IAF is a stable trait, making it useful for within-person longitudinal tracking. By following changes in IAF over time, deviations from normal CNS functioning, such as onset or progression of disease, can be monitored.

## Introduction

The alpha frequency, generally between 8 and 12 Hz, is the dominant frequency of the human electroencephalogram (EEG), ^1 2^ as shown in Figure 1. Measured during relaxed wakefulness, the power of the alpha rhythm is typically strongest in the eyes-closed condition. The peak of the alpha frequency (AF) can provide information on central nervous system functioning as well as the status of mental health and cognitive functioning. Significant correlations between AF and a large variety of cognitive measures have been observed, ^3 4 5 6 7^ perhaps suggesting links between AF and speed of information processing, the timing of neural inhibition, or the gating of information in the brain. ^8 9 10 11^

**Figure 1.**
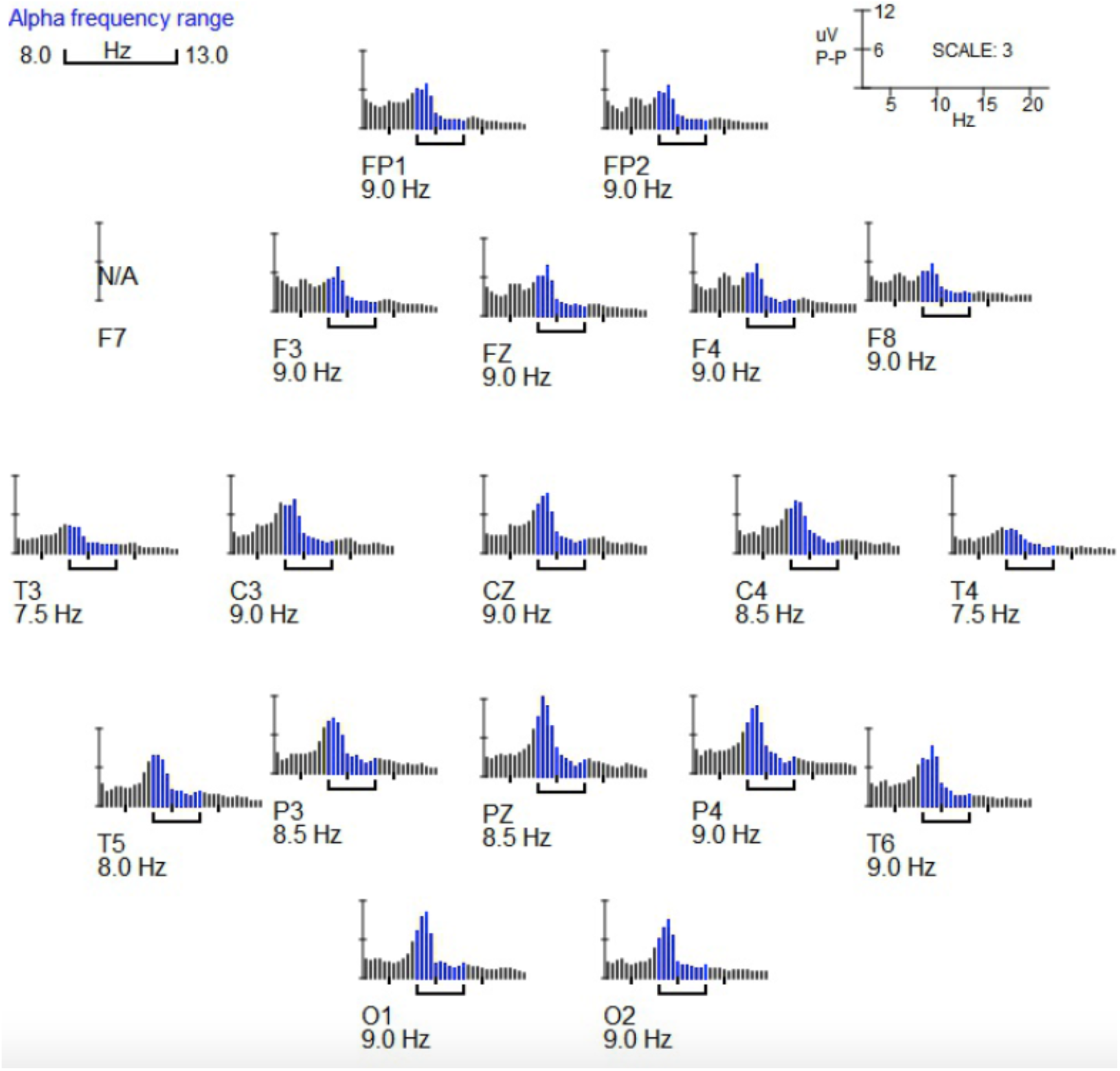
Frequency versus EEG amplitude spectra extracted from the WAVi platform for a sample subject, a healthy 25-year-old control. Here the alpha window is displayed by the brackets below the spectra (set in this case from 8-13Hz to guide the eye). Notice the strong alpha peak at all locations, with more theta contribution in the frontal areas for this patient.

Clear changes in the alpha rhythm are seen as the human brain ages, including AF slowing, reduction in alpha power, and a shift in the posterior-to-anterior direction. ^5 12 13 14 15 16 17 18 19 20 21 22^ Slowing of the EEG has also been found to indicate CNS pathology, in particular slowing of AF has been observed repeatedly in patients with dementia. ^23 24 25 26 27 28 29 30 31 32 33 34^

AF alone, however, may be of limited diagnostic value because the majority of these studies focus on comparing groups of individuals with respect to group differences and this may not always translate to identification of CNS pathology at the individual level. For example, even if the mean AF of a group with the diagnosis of dementia is found to be slower than the mean AF of healthy controls, an individual may still be located within the distribution of healthy adults.

An individual’s peak alpha frequency (IAF), however, is considered a stable marker of a neurophysiological trait and can therefore provide longitudinal information on a patient.^35 36 37 38 39 40 41^ Even when large gains in cognitive performance are achieved, the marker remains stable. ^42^ Accounting for some of this stability is the observation that features of the alpha rhythm are likely to have a strong genetic component. ^43 44 45 46 47 48 49^

Given the high stability in the absence of neuropathological processes, IAF may be a valuable marker for monitoring neuropathological changes within individuals. Where single tests may possess poor sensitivity and/or specificity, by following within-person changes over time IAF can be valuable for monitoring deviations from normal CNS functioning, such as the onset or progression of disease. ^42^ For this reason, IAF may be most valuable in tracking healthy individuals and looking for early changes before perhaps even symptoms arise.

For such monitoring to occur, this well-studied metric needs to move from research settings into standard clinical practice, including standard wellness visits. The objective here is to compare a large AF dataset collected in clinic to published research. In particular, we want to validate a clinical method involving a low cost system designed to maximize information and minimize testing times. This includes concurrent testing of eyes-closed resting EEG protocols with eyes-closed EEG evoked response protocols.^50^ To that end the objective of this paper is to (1) validate the use of eyes-closed P300 to collect eyes-closed resting IAF measures and (2) validate data collected in routine clinical settings against published trends of AF over the lifespan (3) validate the within-person stability of IAF from an in-clinic sample.

## Methods

### Subjects

While the subjects for this study were comprised of 2025 subjects from previous or ongoing studies, it is not intended that they represent a normal control for a general population, rather it is intended to provide a target reference. One of the goals of this study is to compare in--clinic data with historical research to test the validity of large-scale screening. This study includes 3 control groups (13-16 years of age, 17-23, and 81-90) which will anchor the resulting age-matched curves as discussed below. It may be the case that these controls perform differently from a found in a normal population, where 2 of these control groups were taken from elite club, High School, or NCAA athletic teams while those in the oldest age range were volunteers living independently, still interested in brain science, and still interested in their brain performance. Each group will be discussed individually, but because of the suspected other-than-normal performance, we will focus on age trends and refer to this reference group as a target reference, with end points as discussed, rather than a normal reference. It is important to note that male/female differences are not the focus of this paper which is to compare trends to literature to establish a reference target.

All studies were approved by appropriate IRB’s and written informed consent was obtained from the participants before study intake.

#### Ages 8-12

73 subjects aged 8-12 were taken from three previous studies: a study that followed athletes over the course of their sports seasons in Texas and Washington, control subjects measured as part a beta test to explore the outcome of an educational/wellbeing intervention program in a school representing low-income students, ^51^ and minors accompanying WAVi study volunteers discussed below. Of these, 57 were accepted as per the artifacting criteria discussed below.

#### Ages 13-16

This control group comprises 94 subjects from a previous study following athletes aged 13-16 over the course of their sports seasons and at 4 different sites. These subjects are participants in youth soccer and youth basketball representing all players from single teams. To follow the objectives of this study (as well as the above-mentioned studies) which involves real clinical settings, our exclusion criteria are minimal. Therefore the only exclusions are players who had lower than 80% yield as per the artifacting criteria outlined elsewhere. ^52^ Of these subjects, 75 were deemed to have sufficient yield and 66 returned and completed a valid post-season second test which will be used to discuss test-retest variability.

#### Ages 17-23

This second control group is taken from a previous study that followed 364 athletes aged 17-23 over the course of up to 4 sports seasons and at 5 different sites. These subjects are participants in NCAA Div. 1 men’s football (172 players, representing all players from a single team), woman’s soccer (29 NCAA Div. 1, representing all players from a single team), men’s high school football (142 players, representing all seniors from a single team), and semipro men’s ice hockey (20 players, representing all players from a single team). ^9^

In these previous studies, these subjects were controls against which pre-contact, post-concussion and return-to-play groups could be compared. To follow the objectives of this study (as well as the above-mentioned studies) which involves real clinical settings, and because the primary marker being studied is nonspecific, our exclusion criteria are minimal. The “control” group, therefore, is a reference group taken from all players participating on these teams and exclusions are limited to the players who fell asleep during the first-year test and passing the artifact criteria discussed below, leaving a total of 313 players comprising the baseline reference group of Table I. Of these subjects, 70 returned injury free to completed a valid second test, which will be used to discuss test-retest variability.

#### Ages 24-30

138 assessments were included for individuals aged 24-30 who were measured in clinic at baseline, where some were to be tracked over the course of various interventions. Subjects include patients who visited the Boone Heart Institute Colorado for a combined preventative cardiology and EEG/ERP evaluation from June 2014 through June 2017. Only first-time patients receiving an initial evaluation were included in the sample, which was also used for a preventative cardiology study. ^6^ Because this is a target reference study, the exclusion criteria are minimal, the criteria being those who were taking beta-blockers or psychiatric medication and those who had lower than 80% yield on evoked responses due to artifact.

Also included were subjects from Natural Bio Health (NBH) Texas for a first-time preventative wellness exam, evaluated from 2017 through 2018; and a random sample of subjects measured for demonstration purposes at 5 medical conferences.

The remaining subjects were volunteers who were known to or associated with the study team and wanted to become pro-active in their brain health. In general, these reference subjects were well educated and wanted to use WAVi to compare pre-intervention to post-interventions where interventions typically included some form of lifestyle change. To follow the objectives of this study, which involves real clinical settings our exclusion criteria are minimal and all volunteers in this age group were analyzed for the purposes of this study.

#### Ages 31-40

217 individuals aged 31-40 were tracked over the course of various interventions and the above-mentioned clinics, conferences, and volunteers.

#### Ages 41-50

325 individuals aged 31-40 were tracked over the course of various interventions and the above-mentioned clinics, conferences, and volunteers.

#### Ages 51-60

397 individuals aged 31-40 were tracked over the course of various interventions and the above-mentioned clinics, conferences, and volunteers.

#### Ages 61-70

249 individuals aged 31-40 were tracked over the course of various interventions and the above-mentioned clinics, conferences, and volunteers.

#### Ages 71-80

64 individuals aged 31-40 were tracked over the course of various interventions and the above-mentioned clinics, conferences, and volunteers.

#### Ages 81-90

Our third control group comprises 42 people taken as volunteers, discussed above. This group were living independently, had not been diagnosed with dementia, and were by definition a population who had experienced what could be called successful cognitive aging. They provide an end point against the 20-year-old athletes for our target reference. Of these subjects, 26 were deemed to have sufficient yield to be included in the study.

#### Ages 24-85 Test-Retest Study Group

This group comprises 67 tests from 8 volunteers to study test-retest for various ERP and qEEG parameters during eyes-closed audio P300.

#### Ages 24-58 Default Network Study Group

This group comprises 108 total tests taken from 8 volunteer control subjects from the test-retest study above and from 22 subjects collected at a clinical site testing various conditions such as mood, attention, and for baseline wellness. These tests are of the reliability of audio P300 protocol as a default network by comparing the qEEG metrics extracted during the standard eyes-closed qEEG protocol to those extracted during the eyes-closed audio P300 protocol, including the theta/beta ratio, left-right alpha asymmetry, and alpha mean frequency in both a control and a clinical setting.

### EEG acquisition and preprocessing

The WAVi Brain Assessment (WAVi Co., Boulder, CO USA) was used to record an electroencephalogram (EEG) at 250 Hz with the position of the electrodes following the International 10-20 system. Reference electrodes were clipped onto the earlobes. EEG eyes-closed resting parameters were recorded on each patient during a 4-min audio P300 and, for some subjects, during a 4-min eyes-closed resting protocol.

### EEG extraction and analysis

WAVi 9.8.18 research software was used to extract and analyze the EEG data (the results of which are intended to create the target ranges for WAVi Scan 1.0-software). Frequency spectra were extracted from the eyes-closed and P300 EEG protocols using standard Fourier methods (Figure 1). This analysis included automatic artifact rejection that excludes noise from EEG data with higher than acceptable amplitudes while also excluding excessive band frequency activities in the standard EEG bands (Delta, Theta, Alpha, and Beta) on an individual channel basis. Files were also visually inspected to confirm proper noise extraction. In addition, when no clear alpha peaks were found, particularly when the delta and/or theta backgrounds were too large, a null result was given. While not a common occurrence, this null result is more common for frontal alpha than for occipital where alpha is generally more pronounced.

Finally, we extract AF for 3 regions of the scalp: Frontal (average frequency for F3, F4, and Fz); Central-Parietal (C-P, average frequency for C3, C4, Cz, P3, P4, and Pz); and Occipetal (average frequency for O1 and O2).

## Results

### AF During P300 and During Eyes-Closed Resting

In order to maximize the information and minimize collection times, we compare IAF extracted during a 4-min eyes-closed Audio P300 (creating the AF of the P300 group) to those values extracted during standard eyes-closed resting (Table II). To minimize subject bias, the same number of eyes-closed and P300 tests were analyzed for each of the 30 subjects (22 subjects measured in clinic and 8 volunteer controls). From 108 reliable tests (80 in the front), no difference was seen between the two methods. Because we assume no difference in AF between these protocols, and to be consistent with the goal of validating a platform that maximizes information and minimizes testing times, the age-related IAF/AF trends of this study are extracted from this audio P300 protocol.

**Table II.**
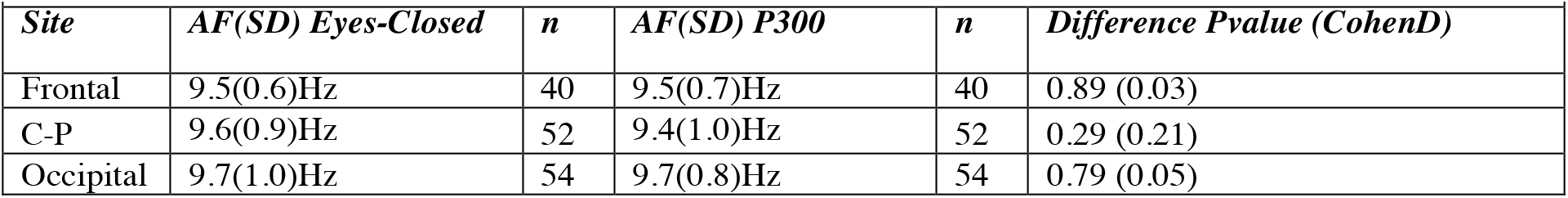
AF extracted from IAF during standard eyes-closed resting and audio P300 protocols.

### AF Age Trends

WAVi in-clinic data for AF over the lifespan, extracted during the eyes-closed P300 protocol, is shown in Figure 1. Strong agreement with previous studies in both magnitudes and age trends is seen. Figures 3-5 show AF over the lifespan for 3 locations of the scalp. As expected, the frequency declines with age but we also see a maturation effect with the fastest frequencies occurring at around 20-30 years of age. From this shape, the best fit for each scalp location is shown and from the resulting curve a target trendline can be established. Note high person-to-person variance where the “fastest” 85-year-olds are still faster than the “slowest” 20-year -olds, reinforcing the utility of IAF for tracking but not always for single tests.

**Figure 2.**
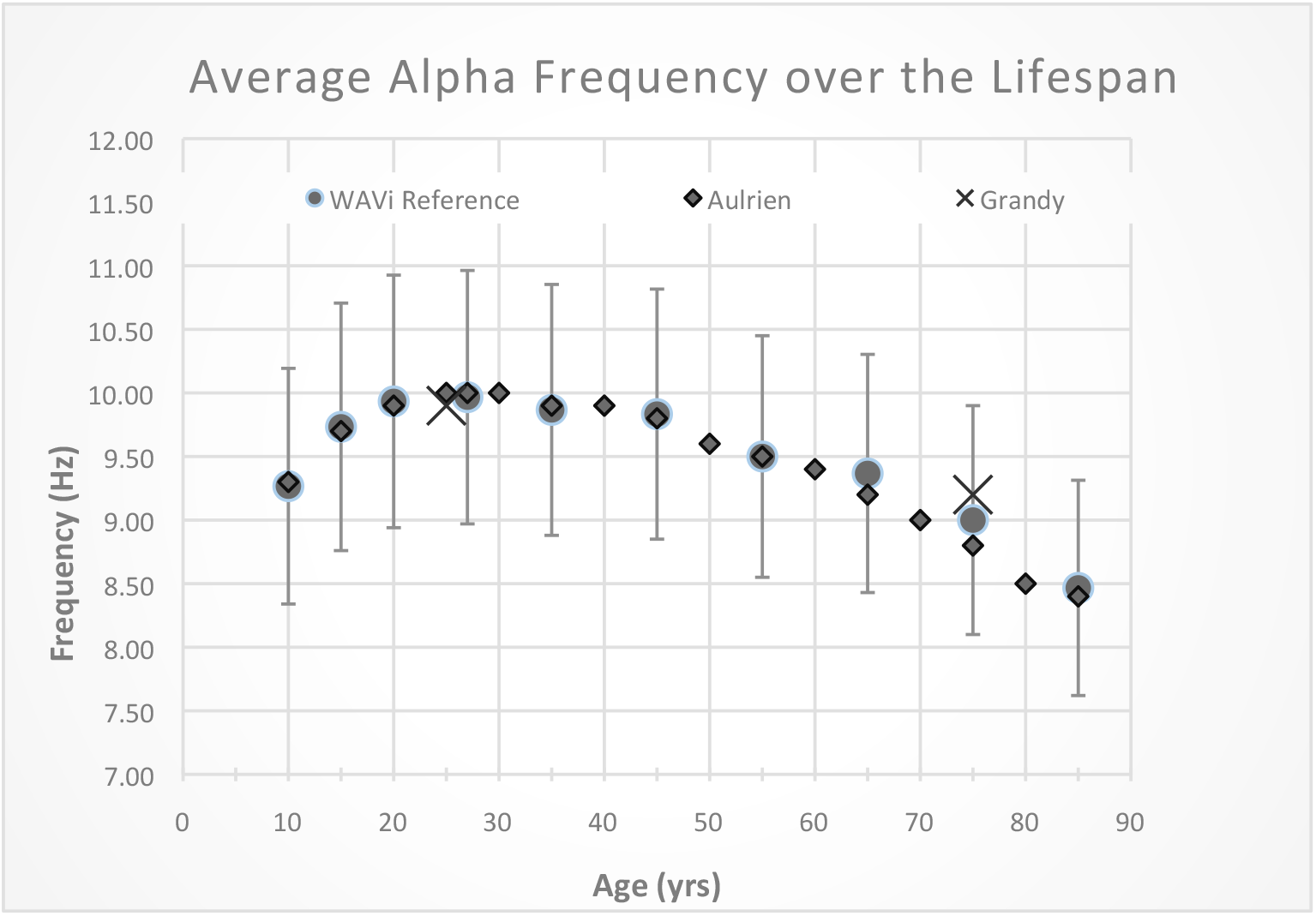
Comparison of Peak Alpha Frequencies extracted in routine clinical settings on the WAVi platform, as a function of age, to various studies. Here we take an average of frontal, central-parietal, and occipital locations.

**Figure 3.**
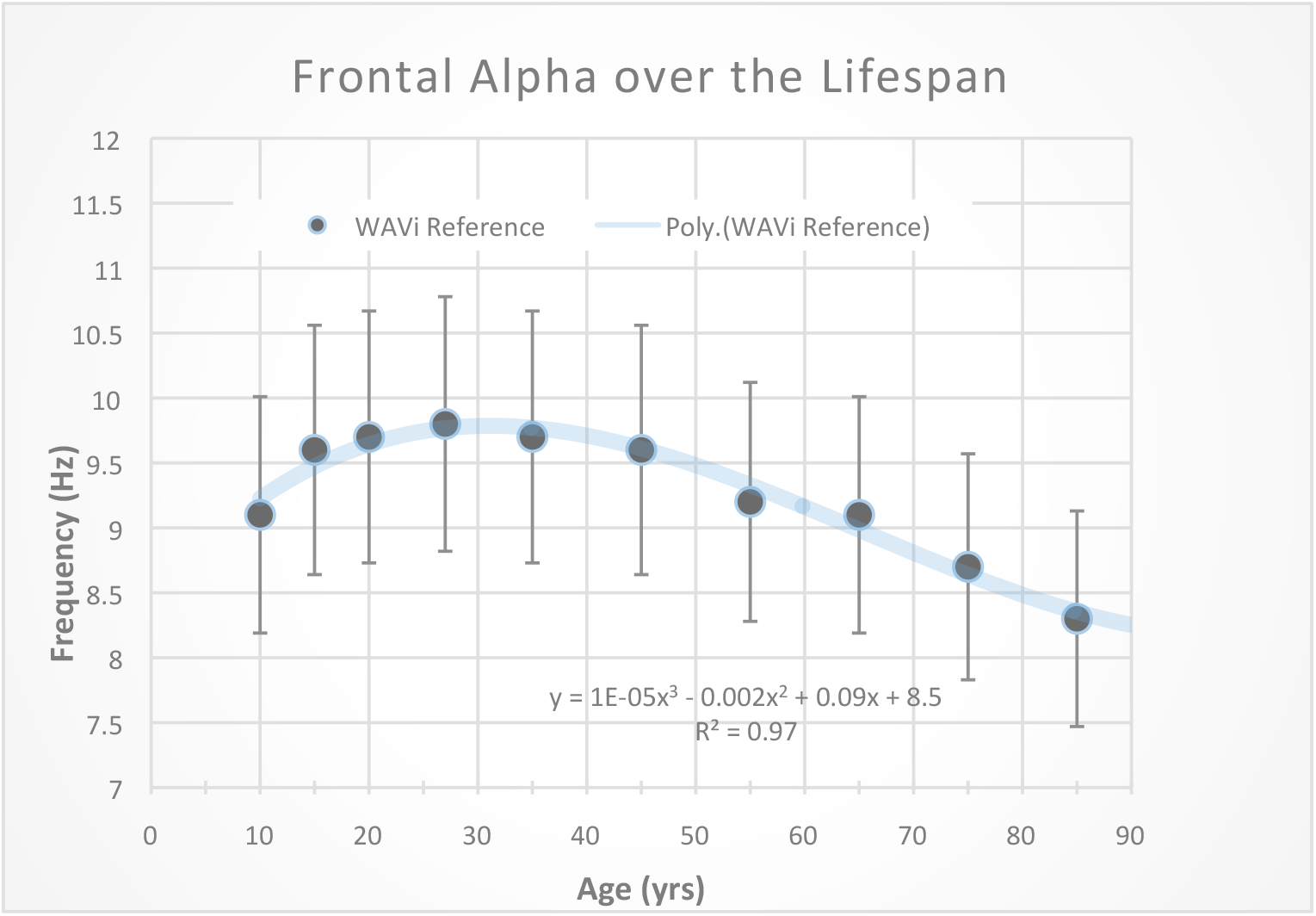
Peak Alpha Frequencies extracted in routine clinical settings on the WAVi platform, as a function of age, for the frontal locations.

**Figure 4.**
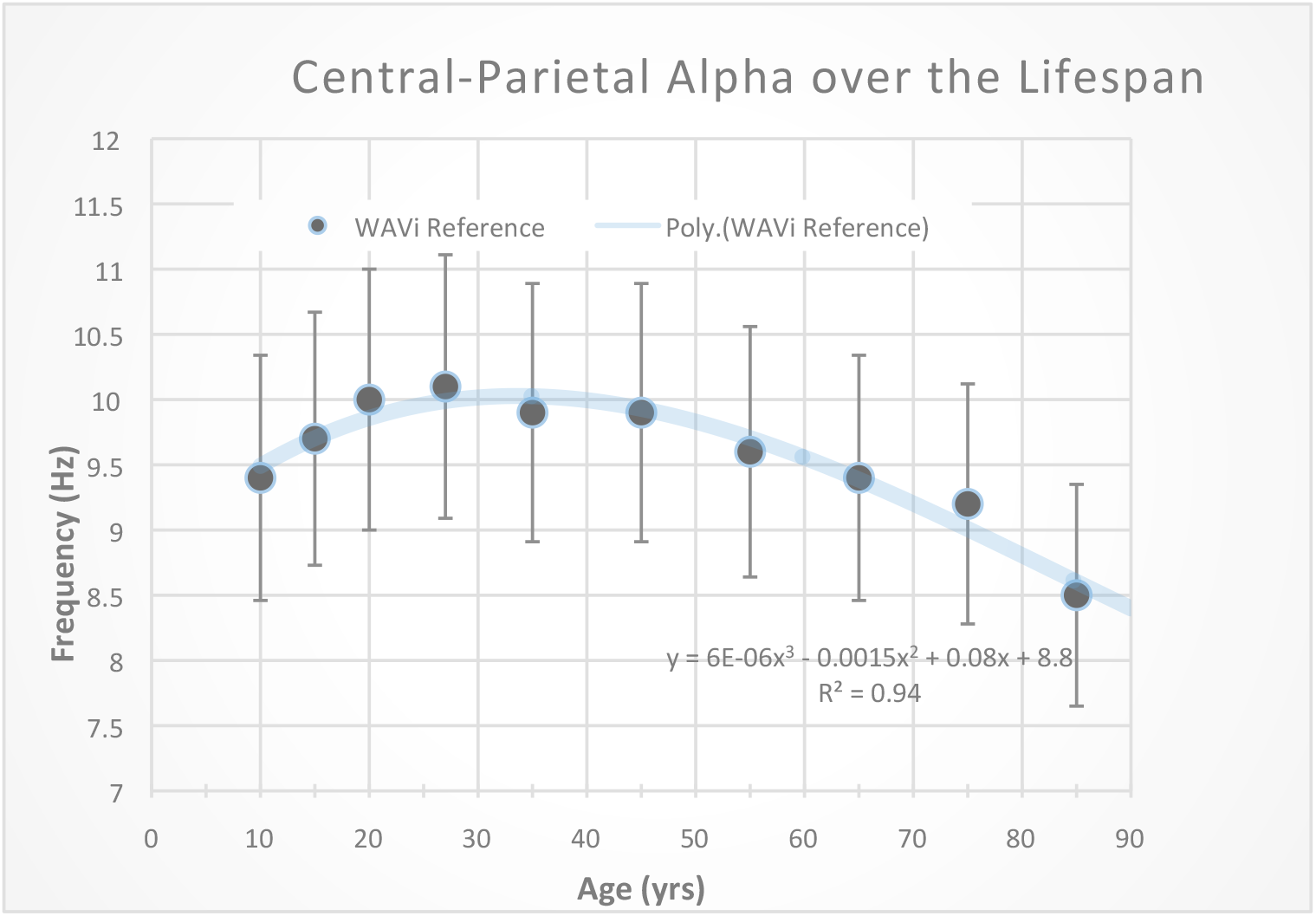
Peak Alpha Frequencies extracted in routine clinical settings on the WAVi platform, as a function of age, for the central-parietal locations.

**Figure 5.**
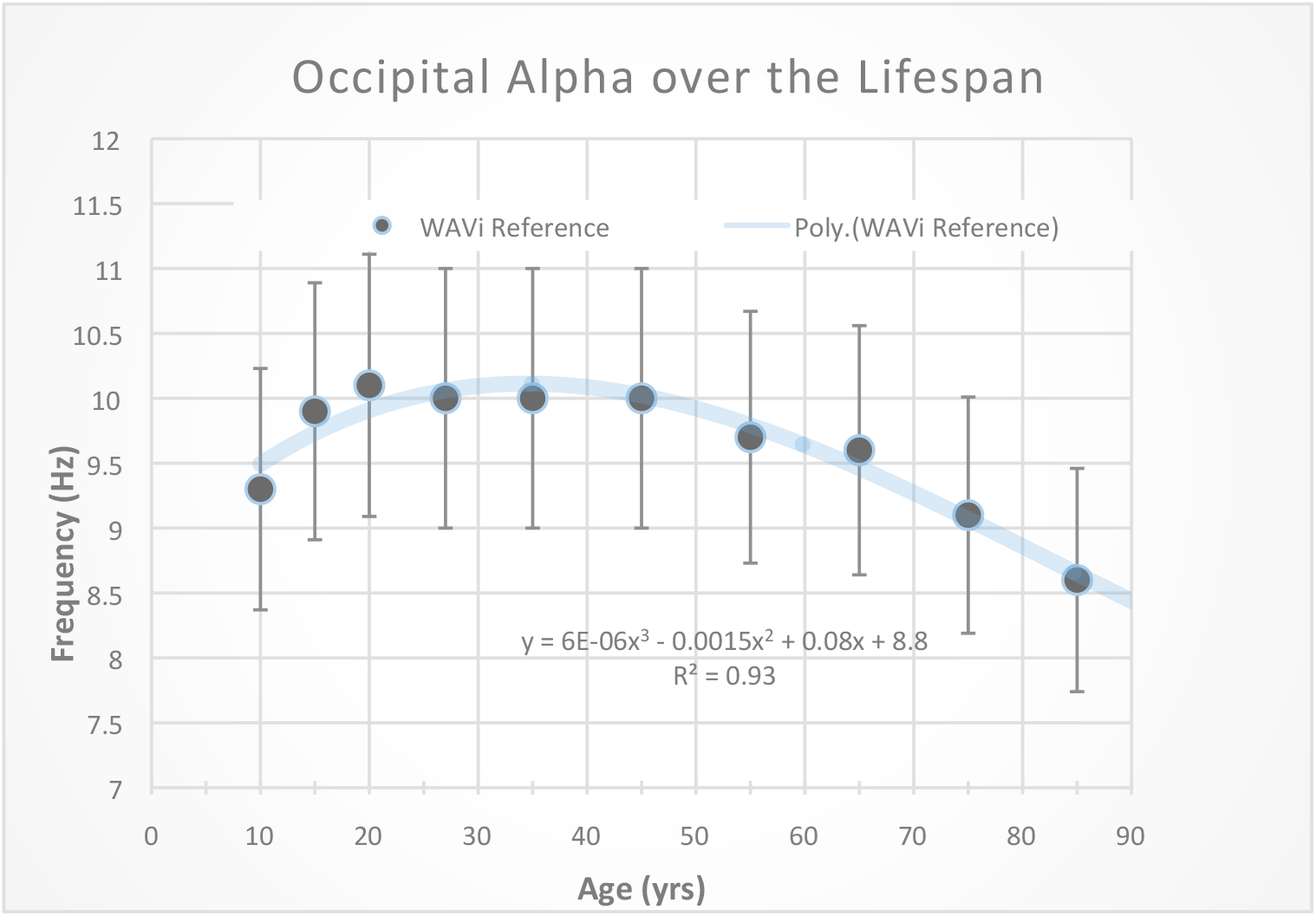
Peak Alpha Frequencies extracted in routine clinical settings on the WAVi platform, as a function of age, for the occipital locations.

A note on variance: It is common practice in medicine to quote “normal ranges” (middle 68 percent of people in a Gaussian distribution) in order to provide context for both the clinician and client. To that end, we have found a variance of +-10% provides a useful target range for all ages and regions.

### Test-Retest Variability

IAF stability over time can be an important characteristic of intact general CNS functioning. While AF values alone may possess poor sensitivity and specificity as a clinical marker, IAF may constitute a robust and easy-to-assess metric for monitoring deviations from normal CNS functioning, such as progression of disease, by following within-person changes in IAF over time. ^42^ As discussed, changes in IAF over time can be an early individual marker of pathology, but such use cases require that IAF remain stable within individuals in the absence of pathology.

Based on these previous studies, the expected intrapersonal variance in IAF from these data is stable. Table III shows test-retest results covering an age range of 24-85 years where 84% of the subjects fell below +-0.2Hz in change from their personal average over the course of 1 month to 2 years. These numbers provide a reference from which longitudinal changes can be studied, either after an event such as concussion, an intervention, or in order to monitor healthy aging.

**Table III.** Expected longitudinal change of IAF, with Coefficients of Variance (CV). Data were extracted over a 1-2 year period from the “test retest group” of Table I.

|  |  |  |
| --- | --- | --- |
| ***Frontal (CV)*** | ***C-P (CV)*** | ***Occipital (CV)*** |
| $\pm 0.2\text{ Hz (3.8)}$ | $\pm 0.2\text{ Hz (2.5)}$ | $\pm 0.2\text{ Hz (3.2)}$ |

Also shown in Table III are coefficients of variation (CV), calculated as 100*SD/Average for each person and then averaged over each group. This provides context to compare IAF to other methods of physiologic assay, say for example HDL measures (CV∼24) and cholesterol (CV∼14). ^50^ Table III CV values are lower than most other clinical assays, reinforcing the trait nature of these metrics and highlighting the clinical utility of longitudinal tracking. ^12^

## Conclusion

In-clinic measures of AF extracted during an eyes-closed auditory P300 protocol corroborate the age-related trends of published research taken over the last several decades. These results also confirm that IAF is a stable trait, making it useful for within-person longitudinal tracking. Regarding IAF collected during P300, it is reasonable to hypothesize that the eyes-closed resting protocol may not be as stable due to lack of control of the state of the patient (i.e drowsiness, concentration, etc.) while the P300 protocol requires a constant cognitive engagement. By following changes in IAF over time, deviations from normal CNS functioning, such as onset or progression of disease, can be monitored.

